# The maturation of infant and toddler visual cortex neural activity and associations with fine motor performance

**DOI:** 10.1101/2024.06.11.598480

**Authors:** Katharina Otten, J. Christopher Edgar, Heather L. Green, Kylie Mol, Marybeth McNamee, Emily S. Kuschner, Mina Kim, Song Liu, Hao Huang, Marisa Nordt, Kerstin Konrad, Yuhan Chen

**Affiliations:** Child Neuropsychology Section, Department of Child and Adolescent Psychiatry, Psychosomatics and Psychotherapy, Faculty of Medicine, RWTH Aachen University, 52074 Aachen, Germany; Department of Psychiatry, Psychotherapy and Psychosomatics, Faculty of Medicine, RWTH Aachen University, Aachen, Germany; Lurie Family Foundations MEG Imaging Center, Dept. of Radiology, Children’s Hospital of Philadelphia, Philadelphia, PA, 19104, USA; Department of Radiology, Perelman School of Medicine, University of Pennsylvania, Philadelphia, PA, 19104, USA; Department of Psychiatry, Perelman School of Medicine, University of Pennsylvania, Philadelphia, PA, 19104, USA; JARA-Brain Institute II, Molecular Neuroscience and Neuroimaging (INM-11), RWTH Aachen & Research Centre Jülich, 52428 Jülich, Germany

**Author notes:** <u>Address for Correspondence: Katharina Otten</u>, Department of Child and Adolescent Psychiatry, Psychosomatics and Psychotherapy Faculty of Medicine, RWTH Aachen University Neuenhofer Weg 21 52074 Aachen, Germany.

**Keywords:** Infants, Magnetoencephalography (MEG), Visual evoked responses (VER), maturation, fine motor, developmental trajectory

## Abstract

Our understanding of how visual cortex neural processes mature during infancy and toddlerhood is limited. Using magnetoencephalography (MEG), the present study investigated the development of visual evoked responses (VERs) in both cross-sectional and longitudinal samples of infants and toddlers 2 months to 3 years. Brain space analyses focused on N1m and P1m latency, as well as the N1m-to-P1m amplitude. Associations between VER measures and developmental quotient (DQ) scores in the cognitive/visual and fine motor domains were also examined. Results showed a nonlinear decrease in N1m and P1m latency as a function of age, characterized by rapid changes followed by slower progression, with the N1m latency plateauing at 6-7 months and the P1m latency plateauing at 8-9 months. The N1m-to-P1m amplitude also exhibited a non-linear decrease, with strong responses observed in younger infants (∼2-3 months) and then a gradual decline. Associations between N1m and P1m latency and fine motor DQ scores were observed, suggesting that infants with faster visual processing may be better equipped to perform fine motor tasks. The present findings advance our understanding of the maturation of the infant visual system and highlight the relationship between the maturation of visual system and fine motor skills.

**Highlights:**

- The infant N1m and P1m latency shows a nonlinear decrease.
- N1m latency decreases precede P1m latency decreases.
- N1m-to-P1m amplitude shows a nonlinear decrease, with stronger responses in younger than older infants.
- N1m and P1m latency are associated with fine motor DQ.

## 1 Introduction

A remarkable process during human development is the rapid change in neural circuit activity during the first years of life. A sensory system that matures relatively rapidly is the visual system (Kandel,2013;Richards & Conte,2020). The visual system provides a means of exploring the world, playing a crucial role in early development (Wallace et al.,2004). For example, visual input is important for the development of fine-motor and locomotor skills (Niechwiej-Szwedo et al.,2017), as well as facilitating phoneme discrimination by providing information about speech articulation (Teinonen et al.,2008).

Assessment of visual evoked responses (VERs; also referred to as visual evoked potentials in electroencephalography(EEG) and visual evoked fields in magnetoencephalography(MEG)) provides a non-invasive way to study the visual system. VERs are electrophysiological signals generated in the visual cortex in response to visual stimuli (Mahajan & McArthur,2012). VERs can be quickly obtained using a checkerboard pattern (Taylor & McCulloch,1992). In adults, the VER to a checkerboard pattern is characterized by a negative peak (N1 or N1m, the magnetic analogous of N1 response), followed by a positive peak (P1/P1m), followed by a negative deflection (N2/N2m)(Thompson et al.,2024). The present study focused on the P1m, this the most commonly VER. Given the International Society of Clinical Electrophysiology of Vision (ISCEV) recommendation to measure P1m amplitude from the proceeding N1m peak (Odom et al.,2016), the N1m response was also examined.

As detailed in the following paragraphs, studies on the development of P1/P1m latency and amplitude have provided mixed results, and studies on the development of N1/N1m during the first two years of life are scarce. In the present study, a cross-sectional sample was recruited to assess the development of P1m and N1m from 2 months to 3 years of age, with longitudinal measures obtained from a subset to examine between-infant variability in the maturation of VERs.

### 1.1 VER Latency and Amplitude

VERs are used to assess the maturation of the visual system, with the P1/P1m the most consistently observed visual component in young children (Creel et al., 2019). EEG studies report rapid maturation of the P1 latency, with an adult-like latency of ∼100 ms observed between 6-12 months (Crognale et al.,1993;Kos-Pietro et al.,1997;McCulloch et al.,1999;McCulloch & Skarf,1991;Taylor & McCulloch,1992). Some studies, however, suggest continued maturation of the P1m after infancy (Ellingson et al.,1972;Emmerson-Hanover et al.,1994). Only a few studies have reported on the infant visual N1/N1m response. Whereas some studies report the emergence of the N1 at 2-4 months (Morrone et al.,1996;Taylor & McCulloch,1992), Hammarrenger et al. (2003) report the appearance of the N1 slightly later, at 6-8 months.

Evoked response amplitude measures reflect the cumulative activity of excitatory and inhibitory postsynaptic potentials (Purpura,1959;Siper et al.,2016), with the amplitude changing as a function of myelination (Heidari et al.,2019;Jones & Brusa,2003), synaptogenesis, and neural pruning(see example from the auditory literature: Moore & Linthicum,2007). The few EEG N1 studies have observed a response between 2-6 months (Hammarrenger et al.,2003;Morrone et al.,1996;Taylor & McCulloch,1992). Whereas some EEG studies report no age-related P1 amplitude change in response to pattern reversal stimuli (Jiang et al.,2023;Lenassi et al.,2008), other studies report a gradual increase in P1 amplitude until ∼2 years old (Kos-Pietro et al.,1997), and some a maximum P1 amplitude around 2-6 months of age (Crognale et al.,1993; Hammarrenger et al.,2003; Jensen et al.,2019; Lippé et al.,2007). The inconsistent P1 amplitude findings are likely due, in part, to the fact that some studies measure P1 amplitude as a baseline-to-peak measure (Hammarrenger et al.,2003;Lippé et al.,2007;Tremblay et al.,2014) and some P1 amplitude from the peak of the previous N1 (Crognale et al.,1993;Jensen et al.,2019;Odom et al.,2016).

### 1.2 VERs: Associations with Behavior and Clinical Conditions

Kim et al. (2018) found that children under 23 months of age with a delayed EEG P1 latency (defined as > 115 ms) tended to have low scores on the psychomotor development index(PDI) and the mental development index(MDI) of the Bayley Scales of Infant and Toddler Development (Bayley-2). Torres-Espínola et al. (2018) showed that a delayed P1 latency at 3 months was associated with a lower cognitive composite score(Bayley-3) at 18 months in infants born to mothers who were overweight, obese, or had gestational diabetes. Jensen et al. (2019) reported a negative association between P1 latency measures obtained at 6 months and the Mullen Scales of Early Learning(MSEL) visual reception and fine motor domain scores obtained at 27 months in infants growing up in a Bangladesh urban slum.

Using MEG and a prosaccade task, Pervin et al. (2022) showed a later latency to peripheral visual stimuli and a reduced amplitude to central stimuli in fetal alcohol syndrome versus TD children. The above findings suggest that during development an earlier P1/P1m latency indicates a more mature visual system, and that P1/P1m latency predicts future cognitive and motor ability.

Recent studies suggest that the P1 amplitude provides valuable information. Siper et al. (2022;2016) found that children with autism spectrum disorder and children with Phelan-McDermid syndrome had a smaller N1-to-P1 amplitude than healthy controls, which the authors suggested reflects an excitatory/inhibitory imbalance. Saby et al. (2023) observed negative associations between N1 and N1-P1 amplitude and clinical symptom severity in children with Rett syndrome. Jensen et al. (2019) showed that a stronger P1 amplitude at 6 months was associated with a higher 6-month MSEL cognitive score, and that the P1 amplitude in a cohort of 36-month-old preschoolers was associated with FSIQ scores at 60 months old. Overall, studies indicate that VER latency and amplitude are clinically informative.

### 1.3 Study Aims

Left and right visual cortex responses in a cross-sectional and longitudinal subsample of infants and toddlers 2 months to 3 years old were obtained. Study objectives were: (1) to determine the developmental trajectory of left and right hemisphere N1m and P1m latency and left and right hemisphere N1m-to-P1m amplitude; and (2) to determine if latency and amplitude measures predict performance on assessments examining cognitive or fine motor skills.

## 2 Materials and Methods

### 2.1 Participants

This study was approved by the Institutional Review Board at the Children’s Hospital of Philadelphia. All families gave written informed consent. Inclusion criteria were as follows: (1) no history of seizure disorder and no first-degree relative with a seizure disorder; (2) no premature birth (< 37 weeks gestation); (3) no non-removable metal in the body; (4) no known hearing or visual impairment (as indicated by passing newborn hearing and vision screening); and (5) no concerns regarding developmental delay (based on parent-report questionnaires, medical records, and scoring within 2 standard deviation of the mean in the developmental assessments described below).

The cohort included infants and toddlers participating in three different studies using the same MEG system and the same visual task. MEG data during a visual task was obtained from 140 infants, 46 of which had longitudinal data (only 1 of the 3 studies collected longitudinal data). To construct a cross-sectional sample, for the children with longitudinal data, one dataset per infant was randomly selected. Of the 140 infants included in the cross-sectional sample, 132 had evaluable data. MEG data from eight infants was excluded for the following reasons: crying during the scan or not watching the screen (N = 3), poor data quality due to excessive movement or magnetic noise artifacts (N = 5). Of the 46 infants in the longitudinal sample, data were obtained from 32 infants at 2-6 months (Time 1), 29 infants at 7-11 months (Time 2), 28 infants at 12-23 months (Time 3), and 23 infants at 24-36 months (Time 4) (see Table 2). Among these, 26 infants had data available from two time points, 17 from three time points, and 3 from four time points. Of the 46 longitudinal infants, 2 were excluded due to unevaluable MEG data, thus resulting in an evaluable longitudinal sample of 44 infants.

### 2.2 Developmental Milestone Measures

Developmental milestones were assessed via parent report using the Vineland Adaptive Behavior Scales (VABS-3; Sparrow et al., 2016). Depending on the study, direct clinical observation measures were obtained using either the Bayley Scales of Infant and Toddler Development - Third Edition (Bayley-3; Bayley, 2006), - Fourth Edition (Bayley-4; Bayley & Aylward, 2019), or the Mullen Scales of Early Learning (MSEL; Mullen, 1995). The VABS-3 provides adaptive behavior composite standard scores (ABC score) based on three subdomains: communication, daily living skills, and socialization. The Bayley-3/Bayley-4 assesses developmental function in children from 1-42 months in 5 domains: cognition, motor, language, socio-emotion, and adaptive behavior. The MSEL assesses developmental function in children from birth to 68 months in 5 domains: gross motor, visual reception, fine motor, expressive language, and receptive language.

As studies have shown associations between VERs and cognitive function and fine motor ability (Jensen et al., 2019; Kim et al., 2018; Torres-Espínola et al., 2018), the examined developmental milestone measures included cognitive function and fine motor ability. Performance was measured by estimating an age-independent developmental quotient (DQ) for the cognitive/visual and fine motor domain. Scores from the Bayley-3/Bayley-4 and MSEL were combined, given that they demonstrated strong correlations (Lense et al., 2014). For the cognitive/visual DQ, scores from the MSEL visual reception domain or the Bayley-3/Bayley-4 cognitive subscale were used. For the fine motor DQ, scores from the MSEL fine motor domain or Bayley-3/Bayley-4 fine motor domain scores were used (Torras-Mañá et al., 2016). DQ was calculated by dividing the age-equivalent (AE) score by the chronological age and multiplying the result by 100. Age equivalents were derived from the MSEL or Bayley-3/Bayley-4 standardization sample as the medial raw score for a particular age level. DQ is a ratio of chronological age to developmental age in each domain, indicating how the child is performing a task relative to children of their own age.

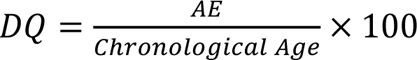

Of note, as one of the longitudinal studies did not perform clinician administered developmental assessments at the first visit (2-7 months) to reduce the burden on families with young infants, developmental data were not available from some of the younger infants (N = 53).

### 2.3 MEG Data Acquisition and Visual Task

MEG data were recorded in a magnetically shielded room (Vacuumschmelze GmbH and Co., KG, Hanau, Germany) using the Artemis 123^TM^ (Tristan Technologies Inc., San Diego, CA, United States) with a sampling rate of 5000 Hz. Artemis 123 was designed for use with children from birth to 3 years of age and features a coil-in-vacuum sensor configuration which minimizes the distance between helmet surface and sensors (Edgar et al., 2015; Roberts et al., 2014). During the MEG recording, infants were presented with a contrast-reversing checkerboard pattern on a screen positioned above their head, with a viewing distance of 55 cm. The pattern was made of 32 x 32 squares and was contrast-reversed at a rate of 1 Hz. The stimulus field subtended a visual angle of 10° x 10°, and the background luminance was set to approximately 50 cd/m2. The visual task lasted 120 seconds with 120 trials collected.

A fabric cap with 4 head position indicators (HPIs) was placed on the child’s head for continuous head position monitoring during the MEG recording. Before the MEG scan, the child’s head shape, anatomical landmarks (nasion, right and left preauricular points), and locations of the 4 HPI coils were digitized using a FastSCAN System (Polhemus, Colchester, VT). During the MEG exam, a research assistant experienced in scanning infants was present to help the parent keep the child calm and alert. A variety of strategies were used to engage the child during the scan (see details in Chen et al. 2021, 2022, 2023). If the infant was not looking at the visual stimuli or was falling asleep, the task was paused or stopped. MEG data acquisition resumed once the infant was again attending to the visual stimuli.

### 2.4 MEG Data Preprocessing and Magnetic Source Analyses

MEG data were analyzed using Brainstorm (Tadel et al., 2011) (http://neuroimage.usc.edu/brainstorm). The raw MEG data were down-sampled to 1000 Hz, and then band-pass filtered 3-55 Hz (even-order linear phase FIR filter; low transition: 1.5 to 3.0 Hz, high transition: 55-63.25 Hz, stopband attenuation: 60 Hz). Heartbeat artifact was removed via independent component analysis (ICA). Artifacts related to head movement and muscle artifact were removed. Trials with MEG activity exceeding 500 fT, due to excessive motion or magnetic noise artifact, were also removed. Visual evoked responses were obtained by averaging artifact-free epochs 200 ms before stimulus onset to 500 ms post stimulus onset. On average, 102.0 ± 31.2 artifact-free trials were used to obtain the evoked response.

MEG data were co-registered to age-appropriate infant MRI templates, available from 1 to 24 months (O’Reilly et al., 2021; Richards et al., 2016). An affine transformation accommodated global scale differences between the infant’s anatomy and the atlas. Digitized surface points from FastSCAN representing the shape of the infant’s head (>10,000 points) were used to co-register the MEG and structural MRI template. Using the VER, whole-brain activity maps were computed using Minimum Norm Estimates (MNE) with constrained orientation (Hauk, 2004; Hämäläinen & Ilmoniemi, 1994; Lin et al., 2006; Matsuura & Okabe, 1995). Activity was mapped to a ∼15,000 vertice cortical-surface source space as a function of time (one millisecond resolution). For each infant, an MEG noise covariance matrix was obtained from an empty room recording obtained prior to each infant’s scan. MNE solutions were computed with normalization as part of the inverse routine based on the noise covariance.

### 2.5 Infant Visual N1m and P1m Responses

The present study examined the infant N1m latency, P1m latency, and the N1m-to-P1m peak-to-peak amplitude. For each infant, the VER magnetic field topography was examined (positive source (red) and negative sink (blue)) - see Figure 1(b)) to determine the approximate latency of the left and right N1m response and the P1m response (Figure 1(c)). Left and right primary visual cortex regions of interest (ROI) were identified based on each subject’s MNE solution at the time of the P1m response (+/- 20 ms, see Figure 1(d)), with a threshold of 5 nAm current density and a cluster threshold of 30 cortical surface vertices in the left and right hemisphere. Across infants, the P1m consistently localized to the medial aspect of the occipital lobe in the left and right calcarine sulcus (i.e., primary visual areas). Source timecourses were extracted from each infant’s MNE solution via averaging the source timecourses across all vertices within the left and right visual ROI. Figure 1(e) shows left and right primary visual source time courses for representative infants at 3, 6, 9, and 12 months. The left and right P1m peak latency from the source timecourses were identified using in-house software. The left and right N1m latency were identified as the time point with the strongest response preceding the P1m peak showing a reversal of the P1m magnetic field pattern. Left and right amplitude was measured as the N1m-to-P1m amplitude (Odom et al., 2016).

**Figure 1.**
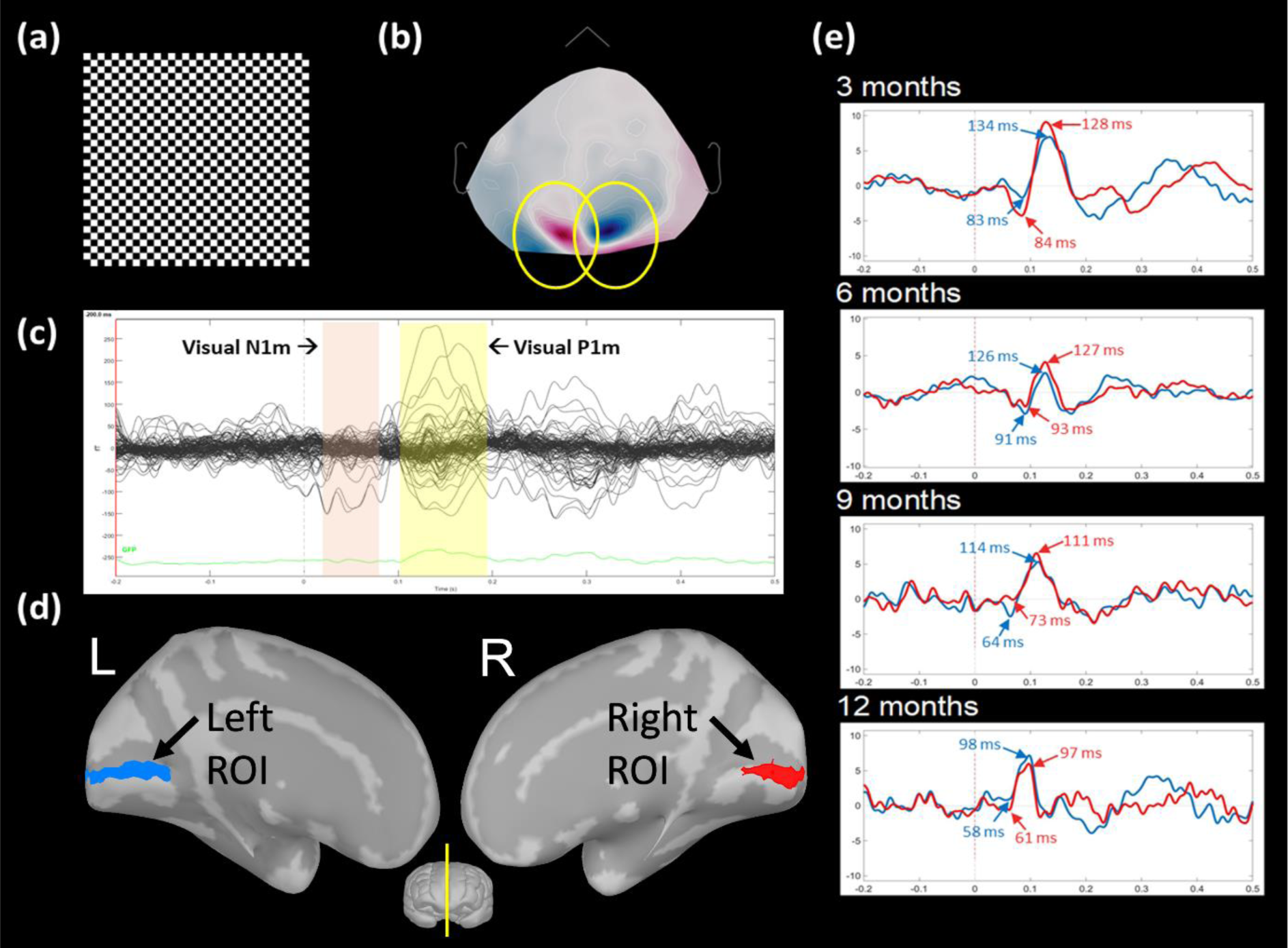
(a) Checkerboard visual stimuli. (b) Example of the sensor magnetic field pattern for the P1m response. (c) Averaged VERs from a representative infant. (d) Left and right visual regions-of-interest (ROIs), denoting visual cortices. (e) Visual source timecourses from 4 representative infants, with right N1m and P1m peak latency shown in red, and left N1m and P1m peak latency shown in blue.

### 2.6 Statistical Analyses

Statistical analyses were conducted using JMP statistical software (JMP Pro, Version 16, SAS Institute Inc., Cary, NC, 1989–2023). Male and female age and developmental milestone scores were compared using two-tailed Student’s t-tests. Analyses were performed separately for the cross-sectional and longitudinal cohorts. Two-way ANOVAs explored the effect of sex and hemisphere on the VER measures. To assess how N1m and P1m latency and N1m-to-P1m amplitude develop as a function of age, the fit of a linear model versus the fit of two non-linear exponential decay models were compared. A LMM examined maturation of the VER in the longitudinal sample, with separate analyses for each VER measure as a dependent variable, and with age (log transformed) as a fixed effect and subject as a random effect. Finally, in the cross-sectional cohort, regressions investigated associations between the VER developmental milestone measures.

## 3 Results

### 3.1 Demographic and Descriptive Statistics

Table 1 shows the demographic information for the cross-sectional cohort (132 infants). There were no differences between females and males for age (*t(*128.8) = 0.59, *p* = 0.56), VABS-3 ABC standard score (*t*(79.80) = -1.60, *p* = 0.11), the cognitive/visual DQ (*t*(63.38) = 0.36, *p* = 0.72), or the fine motor DQ (*t*(61.34) = 0.04, *p* = 0.97). Table 2 shows the demographic information for the longitudinal cohort (44 infants). In the longitudinal cohort females and males also did not differ in age, VABS-3 ABC standard score, cognitive/visual DQ, or fine motor DQ at any time point.

**Table 1.** Demographics: cross-sectional sample.

|  | Female | Male | All |
| --- | --- | --- | --- |
| **Age (months)** | (N = 56) | (N = 76) | (N = 132) |
| Mean (SD) | 9.9 (7.1) | 10.7 (8.3) | 10.4 (7.8) |
| **VABS-3 ABC score** | (N = 37) | (N = 56) | (N = 93) |
| Mean (SD) | 96.5 (9.8) | 93.3 (9.8) | 94.6 (9.9) |
| **cognitive/visual DQ** | (N = 31) | (N = 48) | (N = 79) |
| Mean (SD) | 103.9 (15.0) | 108.7 (21.1) | 106.8 (19.0) |
| **fine motor DQ** |  |  |  |
| Mean (SD) | 100.4 (17.5) | 102.3 (17.8) | 101.6 (17.6) |

**Table 2.** Demographics: longitudinal sample.

|  | **Time 1**<br>2-6 months |  | **Time 2**<br>7-11 months |  | **Time 3**<br>12-23 months |  | **Time 4**<br>24-36 months |  |
| --- | --- | --- | --- | --- | --- | --- | --- | --- |
|  | Female | Male | Female | Male | Female | Male | Female | Male |
| **Age (months)** | (N = 18) | (N = 14) | (N = 11) | (N = 16) | (N = 11) | (N = 17) | (N = 9) | (N = 14) |
| Mean (SD) | 4.6<br>(1.4) | 5.1<br>(1.3) | 9.6<br>(1.3) | 9.6<br>(1.7) | 17.9<br>(2.3) | 18.9<br>(2.5) | 27.6<br>(4.8) | 29.9<br>(5.3) |
| **VABS-3 ABC score** | (N = 7) | (N = 7) | (N = 11) | (N = 15) | (N = 10) | (N = 17) | (N = 7) | (N = 11) |
| Mean (SD) | 92.9<br>(5.7) | 94.1<br>(10.3) | 99.2<br>(7.0) | 98.7<br>(7.5) | 94.5<br>(5.8) | 93.6<br>(7.9) | 93.7<br>(11.5) | 93.9<br>(8.1) |
| **cognitive/visual DQ** | na | na | (N = 11) | (N = 12) | (N = 10) | (N = 15) | (N = 6) | (N = 12) |
| Mean (SD) |  |  | 109.3<br>(20.8) | 112.2<br>(35.5) | 96.8<br>(22.8) | 107.6<br>(17.4) | 112.7<br>(16.1) | 116.2<br>(22.8) |
| **fine motor DQ** |  |  |  |  |  |  |  |  |
| Mean (SD) | na | na | 112.4<br>(23.0) | 115.8<br>(34.6) | 95.3<br>(22.0) | 103.4<br>(11.0) | 104.1<br>(11.8) | 100.5<br>(9.4) |

### 3.2 Infant VERs

Two-way ANOVAs evaluated the effect of sex and hemisphere on the VER measures. As there were no significant interactions with sex or hemisphere for N1m latency (F(1,262) = 0.079, *p* = 0.78), P1m latency (F(1,262) = 0.001, *p* = 0.98) or N1m-to-P1m amplitude (F(1,262) = 0.05, *p* = 0.82), the analyses were re-run with the interaction term removed. No significant main effects were observed for sex (N1m: F(1,263) = 1.3, *p* = 0.25; P1m: F(1,263) = 1.44, *p* = 0.23; N1m-to-P1m amplitude: F(1,263) = 0.26, *p* = 0.61) or hemisphere (N1m: F(1,263) = 0.004, *p* = 0.95; P1m: F(1,263) = 0.001, *p* = 0.98 ; N1m-to-P1m amplitude: F(1,263) = 0.008, *p* = 0.93). Table 3 reports mean and standard deviation values for the left and right N1m and P1m latency and N1m-to-P1m amplitude for the total sample (final row) as well as average values in 3-month age bins. Given no effects of hemisphere or sex, in the following analyses left and right latency and amplitude measures were averaged, and sex was not included as a factor.

**Table 3.**
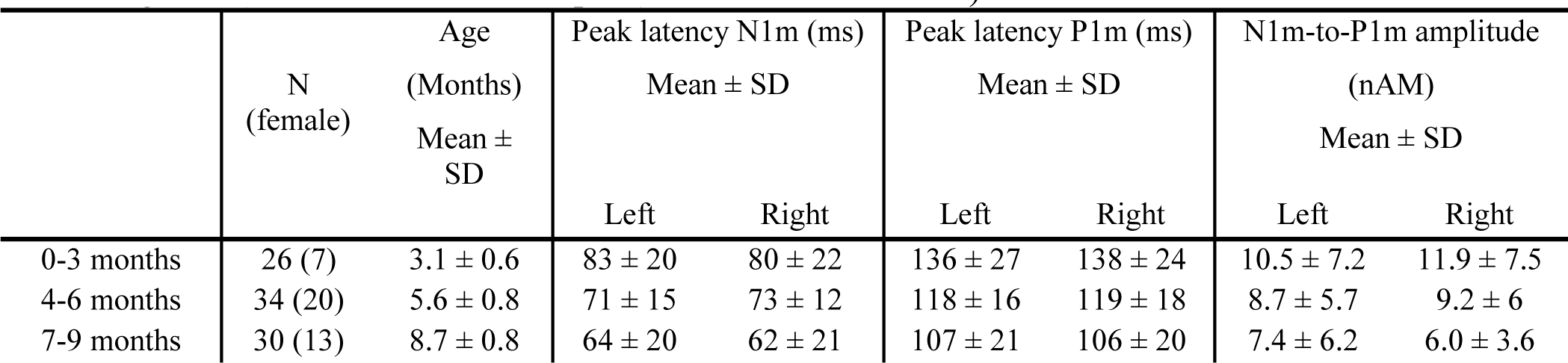
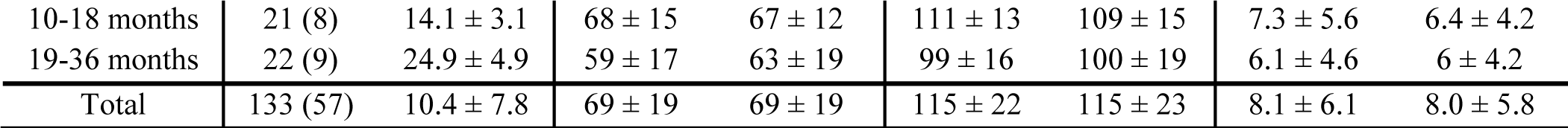
Average right and left N1m and P1m latency and N1m-to-P1m amplitude at different ages (3 month age bins) and for the total sample (cross-sectional cohort)

### 3.3 Maturation Trajectory of Infant VER Measures

The N1m model approximated an asymptotic latency value of 65 ms (95% CI = [61.0, 68.4]), and the P1m model approximated an asymptotic latency value of 104 ms (95% CI = [99.4, 109.0]). Latency decreased rapidly in the first few months of life and then thereafter decreased more slowly. Table 4 shows predicted N1m latency, P1m latency and N1m-to-P1m amplitude from 2-10 months. Predictions indicate that the monthly latency decrease dropped below 1 ms per month for N1m by 7 months of age and for P1m by 9 months of age (Table 4), indicating stabilization of the latency measures at 7 and 9 months, respectively. The N1m-to-P1m response was strongest in infants below 3 months. The relationship with age was best captured by a nonlinear exponential decay model with 3P (N1m-to-P1m amplitude: Linear Fit: AIC = 803.82; Exponential 2P: AIC = 802.16; Exponential 3P: AIC = 798.41, *R*^2^ = 0.12, RMSE = 4.88, BIC = 809.63). The model estimated an asymptotic value of 5.9 nAm (CI = [4.1, 7.8]). To investigate the maturation of the N1m latency, P1m latency and N1m-to-P1m amplitude in the longitudinal sample, LMMs were run with the VER measure as the dependent variable, log-transformed age as a fixed effect, and subject entered as a random effect. Negative associations between age and all three VER measures were observed indicating that the N1m latency (B = -7.84, SE = 2.06, t(92.8) = -3.81, *p* < 0.001), the P1m latency (B = -13.49, SE = 1.95, *t*(85.6) = -6.92, *p* < 0.001), and the N1m-to-P1m amplitude (B = -1.82, SE = 0.53, t(99.9) = -3.41, *p* < 0.001) decreased with age. Figure 2 illustrates the non-linear maturation of the N1m latency (top row), P1m latency (middle row), and N1m-to-P1m amplitude (bottom row) in the cross-sectional and longitudinal cohorts.

**Figure 2.**
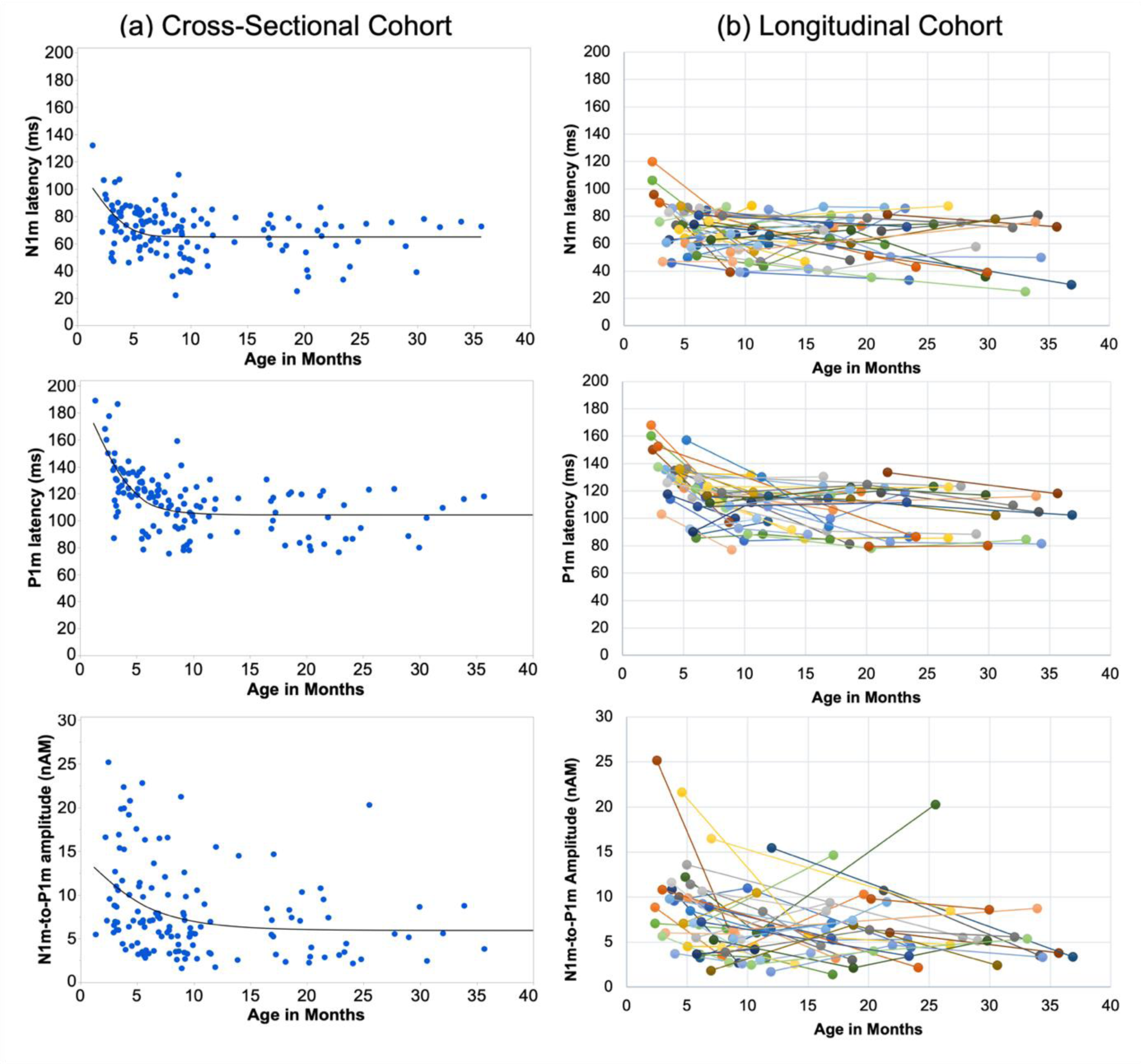
The non-linear maturation of the N1m latency (top row), P1m latency (middle row) and the N1m-to-P1m amplitude (bottom row) as a function of age in the cross-sectional cohort (a) and the longitudinal cohort (b). In the all plots, left and right hemisphere values were averaged. In the panels to the right, each color depicts an individual child.

**Table 4.** Predictions for N1m and P1m latency (ms) and N1m-to-P1m amplitude (nAm) for infants 2-10 months, based on fitted non-linear exponential decay models (3P).

|  | N1m<br>latency | P1m<br>latency | N1m-to-P1m<br>amplitude |
| --- | --- | --- | --- |
| 2 months | 98 | 161 | 12.4 |
| 3 months | 82 | 140 | 11.1 |
| 4 months | 74 | 127 | 10.0 |
| 5 months | 69 | 119 | 9.2 |
| 6 months | 67 | 113 | 8.5 |
| 7 months | 66 | 110 | 8.0 |
| 8 months | 65 | 108 | 7.6 |
| 9 months | 65 | 107 | 7.3 |
| 10 months | 65 | 106 | 7.0 |

### 3.4 Associations Between VER Measures and Developmental Milestones

Regression analyses tested if the VER measures were related to cognitive/visual and fine motor development. As previously noted, as no hemisphere difference was observed for any VER measure, an averaged VER measure was used. No association was observed between any of the three VER measures and cognitive/visual DQ (Figure 3). Associations were observed between N1m latency and fine motor DQ (N1m: *R*^2^ = 0.12, *F*(1, 77) = 10.98, *p* = 0.001), as well as between P1m latency and fine motor DQ (P1m: *R*^2^ = 0.07, *F*(1, 77) = 5.42, *p* = 0.02). Specifically, an earlier N1m and P1m latency was associated with a higher fine motor DQ score. No association was observed between the N1m-to-P1m amplitude and the fine motor DQ score (see Figure 3).

**Figure 3.**
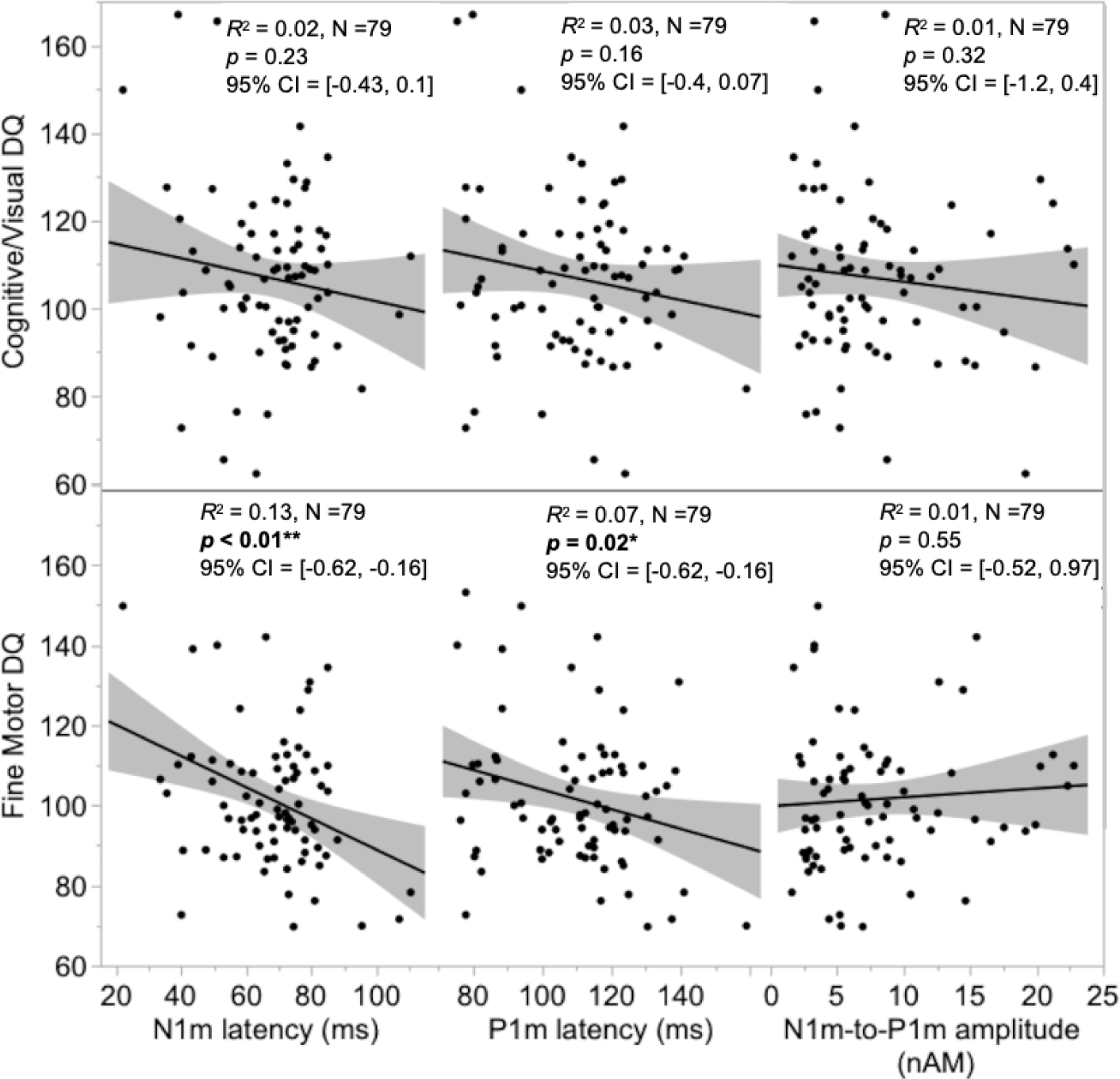
Associations between N1m latency, P1m latency and the N1m-to-P1m amplitude with the cognitive/visual DQ (top) and fine motor DQ (bottom) in the cross-sectional cohort. Significant associations were observed between N1m and P1m latency and the fine motor DQ score.

## 4 Discussion

Via a cross-sectional and longitudinal study design, the present study characterized the maturation of visual cortex neural activity from 2 months to 3 years. Nonlinear maturation of N1m and P1m was observed, with rapid changes in latency and amplitude during the first few months, followed by slower changes thereafter. The latency measures were observed to be of clinical interest, with an earlier N1m and P1m latency associated with a higher fine motor DQ score. The following text discusses the study results with respect to the findings from previous studies.

### 4.1 N1m and P1m latency

Analysis of the cross-sectional and longitudinal samples showed a nonlinear decrease in P1m latency with age, consistent with previous studies (Crognale et al., 1993; Ellingson et al., 1972; Kos-Pietro et al., 1997; McCulloch et al., 1999; Taylor & McCulloch, 1992). Maturation was characterized by rapid changes from birth to ∼8-9 months, followed by a slower change thereafter. N1m latency also decreased nonlinearly with age. As shown in Figure 2, N1m maturation preceded P1m by approximately one to two months, with N1m latency reaching a plateau 6-7 months of age. Present findings contrast with the cross-sectional infant study reported by Hammarrenger et al. (2007) where a parallel latency decrease for N1 and P1 was observed. Compared to P1, the N1 remains relatively understudied, and prior N1 findings are mixed. In the present study, N1m responses were observed in the youngest infants (∼2-3 months). These observations are similar to those reported by Taylor et al. (1992) and Morrone et al. (1996), with both studies reporting the emergence of the N1 at 2-4 months. The present findings are also in agreement with Tremblay et al. (2014). Assessing N1 and P1 responses to a three-phase-reversing vertical sine wave grating stimuli, they observed N1 responses in each age group (3 months, 6 months, 12 months), and reported an age-related N1 latency decrease. In contrast, when showing vertical achromatic sine wave gratings to 0-52 week old infant, Hammarrenger et al. (2003) found that the EEG N1 response only appeared at 6-8 months.

The N1/N1m and P1/P1m latencies reflect, in part, the time it takes for visual information to travel from the retina to the thalamus to the visual cortex. Whereas the N1m response is thought to reflect the propagation of excitation through V1 cortical layers, the subsequent P1m response is believed to reflect intracortical GABAergic inhibition (Lockhofen & Mulert, 2021; Zemon et al., 1980). VERs are influenced by various factors, including the degree of myelination in the visual pathway, the maturation of synaptic connections, and the efficiency of local- and long-range neural transmission (e.g., see studies associating P1 latency to optic radiation diffusion measures (Caffarra et al., 2021; Dubois et al., 2008; Takemura et al., 2020). Of note is that the maturational changes to optic radiation myelination correspond with the maturation of VERs, with a fully myelinated optic nerve (Kinney & Volpe, 2018; Magoon & Robb, 1981) and adult like latencies visual latencies (Crognale et al., 1993; Kos-Pietro et al., 1997; McCulloch et al., 1999; Taylor & McCulloch, 1992) both observed by 7 months. Of note, however, is a study by Takemura et al. (2020), who found that MRI structural tissue parameters along the optic radiation accounted for 22% of variance in P1m latency, and thus demonstrating that other brain features contribute to P1m latency. Takemura et al. suggested pupil size and retinal illumination, tract length, and cortical feedback as other possible factors affecting P1m latency.

### 4.2 N1m-to-P1m amplitude

As detailed in the Introduction, findings regarding the maturation of the N1m-to-P1m amplitude have been inconsistent. In the present study, N1m-to-P1m amplitudes decreased non-linearly, with strong responses observed in the youngest infants (∼2-3 months), followed by a gradual decrease in amplitude. Present findings are similar to studies reporting a maximum amplitude at 2-6 months (Crognale et al., 1993; Hammarrenger et al., 2003; Jensen et al., 2019; Lippé et al., 2007). The N1m-to-P1m amplitude reflects the sum of excitatory and inhibitory postsynaptic potentials, this influenced by myelination, synaptogenesis, and dendritic development (Zemon et al., 1980). The early formation of abundant synapses in the visual cortex causes a net increase in synaptic depolarization, and likely constitutes one of the reasons a larger visual response is observed in younger than older infants. The highest density of synapses in the visual cortex occurs at 2-3 months (Huttenlocher & de Courten, 1987; Rakic et al., 1994), coinciding with the age range that the strongest N1m-to-P1m response was observed in the present study.

Hammarenger et al. (2003) proposed that change to the VER amplitude as a function of age may result from faster development of the magnocellular (M) visual pathway (as reflected by P1 amplitude) than the parvocellular (P) visual pathway (as reflected by N1 amplitude). Hammarenger et al. hypothesize a P1 amplitude decrease from birth to 4-6 months due to the maturation of the P-system, this inhibiting activity of the M-system, and this resulting in mutual inhibition between the two systems and thus smaller P1 responses and larger N1 responses. Another factor possibly contributing to the P1m amplitude is attentional control. Attention to visual stimuli results in a larger VER (Di Russo & Spinelli, 1999; Hoshiyama & Kakigi, 2001). It is likely that developmental-related increases in attention also interact with VER amplitude (Johnson, 1990).

Finally, one reason for inconsistent amplitude findings is likely the result of how the measure is obtained. Although the International Society for Clinical Electrophysiology of Vision recommends measuring the amplitude of the P1/P100 from the preceding N1/N75 peak (Odom et al., 2016), some studies compute the amplitude of these two responses as peak to baseline measures (Hammarrenger et al., 2003; Lippé et al., 2007; Tremblay et al., 2014).

In summary, multiple factors contribute to the maturation of the N1m-to-P1m amplitude.

### 4.3 VER Measures and Developmental Milestones Scores

The present study observed associations between N1m and P1m latency and the fine motor DQ score. To our knowledge, no previous study has linked N1/N1m latency to visual/motor or cognitive outcome in infants or children. Studies have reported associations between P1 latency and current or future behavioral outcomes (Jensen et al., 2019; Kim et al., 2018; Torres-Espínola et al., 2018). As an example, using EEG, Kim et al. (2018) found that children under 23 months with a later P1 latency (defined as > 115 ms) had lower DQ scores than children with an earlier P1 latency (< 115 ms).

The relationship between VERs and fine motor DQ scores observed in the present study is hypothesized to be due to the close correspondence between the maturation of the visual and motor systems. Specifically, visual information regarding an object’s appearance, orientation, and location is essential for initiating grasping and reaching behaviors, as well as carrying out fine motor skills such as grasping and turning, enhancing visual object interaction, and facilitating the exploration of objects in 3D space (Soska et al., 2010). Yu and Smith (2012) noted that infants were more likely to learn the names of objects if an object is named while in their direct visual field or if they are grasping the object (Yu & Smith, 2012). Smith (2013) suggested that holding objects increases sustained visual attention by aiding in the stabilization and alignment of the eyes, head, and hand (Smith, 2013). The necessarily linked development of the visual and motor systems likely precludes our ability to make causal claims regarding the directionality of visual and motor associations. Whereas a maturing visual system aids fine motor development, the development of fine motor skills impacts the development of the visual system.

In contrast to previous studies (Jensen et al., 2019; Perry et al., 1976), no relationship was observed between N1-to-P1 amplitude and cognitive/visual or fine motor scores. This may be due to differences in the age of participants across studies. In particular, whereas Jensen et al. (2019) found that N1-P1 amplitude correlated positively with prospective MSEL domain scores at 27 months, there was no relationship with concurrent MSEL domain scores at 6 months. The authors hypothesized that the MSEL does not capture enough variance in performance in young infants. A similar pattern was observed by Chen et al. (2023), with associations between auditory cortex neural measures and language performance only observed after 12 months, this because very few language production behaviors are observed prior to 12 months and thus the between-subject variability in language ability needed to observe associations was not observed until after the first year of life. Given the above, it is possible that associations between VER amplitude and high-level cognitive processes or behavior may be difficult to assess until after the first year of life.

### 4.4 Future Directions

Findings from the present study inform future VER electrophysiology studies, with the finding that the left and right N1m, P1m, and N1m-P1m amplitude measures do not differ between the left and right hemisphere indicating that when using full-field checkboard stimuli it is likely reasonable to obtain a single N1m or P1m latency measure (e.g., averaging across posterior EEG sensors). In contrast, hemisphere differences are observed for higher-order visual processes such as face encoding, with studies demonstrating the need to separately consider left and right hemisphere activity (Chen et al., 2021). Work regarding hemisphere maturation of visual cortex is needed to determine the validity of sensor measures using divided visual field paradigms.

Finally, of note is the between-subject variability in the VER latency and amplitude measures. As an example, the Figure 2b longitudinal plots show that in infants 10 months old the N1m latency values ranged from 40-80ms and P1m latency values from 80-120ms. This normal variability likely accounts for the latency values observed in Table 3, with a later average N1m and P1m latency value in children 10-18 months than children 7-9 months or children 19-36 months. A similarly large range of latency values (and with many younger children having earlier latencies than older children) was observed for auditory cortex M50 responses in neurotypical children 6-8 years old (Edgar et al., 2020). As detailed in Edgar et al. (2020), an understanding of normal variability in evoked responses is needed to inform clinical research assessing deviations from normality. The present infant VER findings indicate that variability in the maturation of visual cortex neural activity occurs in a manner analogous to maturation of behavioral phenotypes observed “by eye.” For example, although most neurotypical children take their first steps between 9-12 months and are walking by 14-15 months, some neurotypical children do not take their first steps until 16-17 months (WHO, 2006).

### 4.5 Limitations

As previously mentioned, the study compiled data across three studies that varied in the administered neurodevelopmental assessments. To make developmental milestone scores comparable, DQ scores were computed for the visual/cognitive domain and the fine motor domain. Second, the sample size of the longitudinal sample was too small to investigate associations between VER measures and DQ scores obtained at later time points. Third, we did not assess single trials, which might have provided insight into the between-subject latency and amplitude variability. Kovarski et al. (2019) demonstrated the value of single-trial analyses by linking reduced amplitudes in ASD patients to increased inter-trial latency.

### 4.6 Conclusions

The present study showed nonlinear maturation of the N1m and the P1m latency and amplitude, with N1m and P1m latency predicting performance on fine motor developmental assessments. Overall, VER measures were found to be a promising neural brain measure for assessing infant brain development and visual function.

## Data and Code Availability Statement

The MEG data for this project (de-identified) may be obtained via a Data Use Agreement with the Children’s Hospital of Philadelphia. Researchers interested in access to the data may contact Dr. Yuhan Chen at. Software for MEG data analysis associated with the current submission is available at https://mne.tools/stable/index.html and https://neuroimage.usc.edu/brainstorm/.

## Acknowledgements

The authors would like to thank the subjects who participated in the studies and John Dell, Rachel Golembski, Erin Huppman, Peter Lam, Shivani Patel, and Na’Kiesha Robinson who helped with data collection.

## Author contribution

**KO** – Formal analysis, Data Curation, Writing-Original Draft, Visualization; **JCE** – Formal analysis, Funding Acquisition, Conceptualization, Supervision, Resources, Writing-Review & Editing; **HLG** – Investigation, Formal Analysis, Writing - Review & Editing; **KM** – Investigation; **MM** – Investigation; **ESK** – Investigation, Writing - Review & Editing; **MK** – Investigation, Writing - Review & Editing; **SL** – Software; **HH** – Funding Acquisition; **MN** – Writing - Review & Editing; **KK** –Writing - Review & Editing, Supervision; **YC** – Investigation, Formal Analysis, Conceptualization, Methodology, Funding Acquisition, Writing – Review & Editing, Data Curation

## Abbreviations

ADHD: Attention-Deficit/Hyperactivity Disorder
AE: Age-Equivalent
ASD: Autism Spectrum Disorder
Bayley: Bayley Scales of Infant and Toddler Development
BMD: Bipolar Mood Disorder
CI: Confidence Interval
DQ: Developmental Quotient
EEG: Electroencephalography
FASD: Fetal Alcohol Spectrum Disorder
HPI: Head Position Indicator
LMM: Linear Mixed Model
MEG: –Magnetoencephalography
MDI: Mental Development Index
MNE: Minimum Norm Estimation
MSEL: Mullen Scales of Early Learning
PDI: Psychomotor Development Index
ROI: Region of Interest
VABS: Vineland Adaptive Behavior Scales
VER: Visual Evoked Response

